# Drivers of recovery and degradation of riverine benthic macroinvertebrate communities: A nationwide analysis of time series

**DOI:** 10.1101/2024.07.05.602183

**Authors:** Christian Schürings, Willem Kaijser, Svenja Gillmann, Kristin Peters, Jens Kiesel, Armin W. Lorenz, Daniel Hering

**Affiliations:** Aquatic Ecology, University of Duisburg-Essen, Essen, Germany; Department of Hydrology and Water Resources Management, Institute of Natural Resource Conservation, CAU Kiel, Germany; Centre for Water and Environmental Research (ZWU), Essen, Germany

## Abstract

In response to the global freshwater biodiversity crisis this study examines the drivers, influencing recovery and degradation in riverine benthic macroinvertebrate communities across Germany. Utilizing the Asymmetric Response Concept (ARC), which posits that species tolerances to stressors, dispersal capacity, and biotic interactions are critical drivers in aquatic ecosystem recovery, we analyzed a comprehensive dataset from 1568 sites, sampled between 2004 and 2022. Our findings indicate that abiotic stress consistently influences ecological status in both recovery and degradation phases. Interspecific competition shows a stronger positive relationship with ecological status improvements during recovery phases than during degradation, underscoring its importance in the recovery process. Additionally, land use intensity has a nuanced impact: catchments with higher proportions of cropland and urban areas are more likely to recover, while forested catchments are more prone to degradation. This study supports the ARC and highlights the complex interplay of biotic and abiotic variables in shaping ecological outcomes, underscoring the importance of integrated management approaches in freshwater conservation and restoration efforts.

## 1. Introduction

In response to the global freshwater biodiversity crisis (Dudgeon et al., 2006; Reid et al., 2019), a multitude of measures has been introduced, including improved wastewater treatment and some local improved habitat conditions. These measures have made some progress in combatting environmental degradation and biodiversity loss (Pharaoh et al., 2023; van Klink et al., 2020). However, the recovery trajectory has slowed down in the last decades, as particularly shown for benthic macroinvertebrates (Haase et al., 2023; Sinclair et al., 2024; Vaughan et al., 2023).

The success of hydromorphological restoration projects in achieving their expected outcomes varies significantly (Kail et al., 2015; Lu et al., 2019). Although detailed monitoring of individual river restoration projects is rare, some studies report positive ecological outcomes (e.g., Feld et al., 2018; Li et al., 2018). Conversely, other studies have found minimal or no improvements beyond the local habitat scale (e.g., Verdonschot et al., 2016; White et al., 2017). This inconsistency highlights that, while wastewater treatment has shown considerable success, hydromorphological restoration frequently does not achieve the desired ecological benefits. Identifying the key factors that determine the success of these restoration efforts remains crucial.

As many restoration projects primarily focus on the reach scale, their efforts may be constrained by stressors originating beyond the river channel, which is underscored by recent studies identifying strong relations between river health and the adjacent land use intensity (Chen et al., 2023; Schürings et al., 2024a). Yet, when restoration measures successfully improve both hydromorphological conditions and water quality, biota has been shown to recover even in previously strongly degraded streams (Gillmann et al., 2023). This suggests that the recovery of ecosystems is mediated by species tolerances to the environmental conditions in the restored sites (Gillmann et al., 2024a). However, the response of biota is often weaker than the level of improved conditions would suggest (Lorenz et al., 2018). Reasons for the delayed or even absent biotic recovery may be related to remaining stressor impacts (Brettschneider et al., 2019), or to biotic constraints such as dispersal capacity (Sundermann et al., 2011; Enss et al., 2024) or biotic interactions including competition for food and space (Lake et al., 2007).

The interplay of various factors in community development under conditions of degradation and restoration is synthesized in the “Asymmetric Response Concept” (ARC) (Vos et al, 2023). This concept identifies three critical filters — dispersal, species tolerance, and biotic interactions — as key elements for degradation and recovery processes, assigning the individual processes to distinct phases. In short, the ARC predicts that tolerance to environmental stressors is most relevant to community assembly in phases of degradation, while dispersal constraints shape community composition in early phases of recovery, and biotic interactions (e.g. competition with already established species) are a main driver of community assembly in maturing recovering communities.

A recent case study by Gillmann et al. (2024a) on a restored catchment provided initial evidence supporting the ARC, highlighting the importance of species’ dispersal capacity and increasing interspecific competition during recovery, as well as a decrease in the present species’ tolerance to organic pollution with increasing recovery. Further studies by Enss et al. (2024) and Gillmann et al. (2024b) emphasized the relevance of dispersal. However, the role of biotic interactions, especially competition, remains unclear beyond the initial insights from Gillmann et al. (2024a), which were limited to one catchment with eleven sampling sites.

To enhance transferability and gain a better understanding of the importance of competition in recovering versus degrading rivers, we utilized a nationwide macroinvertebrate dataset of 1568 sites, each sampled between two and 13 times between the years 2004 and 2022. We related changes in the macroinvertebrate ecological status (Haase et al., 2004) to changes in abiotic stress and competition (measured by trait similarity) and tested other mediating factors.

Specifically, our research aimed to address the following questions:

1. Does the relevance of species tolerance, measured by the relation between abiotic stress and the ecological status, differ between recovery and degradation conditions?
2. Is the relevance of interspecific competition higher in recovering rivers compared to degrading ones?
3. How do additional variables, particularly the adjacent land use, river types, river systems, the date of sampling and the categorization of the river as natural or heavily modified, mediate recovery and degradation potential of rivers?

These research questions guide our investigation into the variables influencing the ecological status and particularly the role of competition and abiotic stress in ecological recovery processes.

## 2. Methods

### 2.1 Biological data

#### 2.1.1 Data basis

We used a nationwide monitoring dataset of 1557 sites sampled between two and 13 times (average 3.5) between 2004 and 2022 (see Schürings et al., 2024 b, c) combined with own time series data on the Boye catchment (Gillmann et al., 2024a). At each of the sites, macroinvertebrate samples were taken with standardized methods, following the ecological status assessment according to the EU Water Framework Directive, using a multi-habitat sampling method (Haase et al., 2004).

#### 2.1.2 Ecological status

As a proxy for species tolerances, the river type (Pottgiesser and Sommerhäueser, 2008) specific Multimetric Index (MMI) was calculated with the online tool PERLODES (https://www.gewaesser-bewertung-berechnung.de/index.php/perlodes-online.html) using species-level taxa lists. The MMI is an amalgamation of several biodiversity-related indices (differing between river types), which was designed to reflect the impact of various stressors including alterations of morphology, hydrology and water quality (Hering et al., 2006). We here used the standardized version of the MMI ranging between 0 (worst) to 1 (best), the main basis for the ecological status calculation under the Water Framework Directive (European Commission, 2000). For simplification reasons, we refer to the Macroinvertebrate Multimetric Index as the ‘ecological status’ in the following.

#### 2.1.3 Interspecific competition

As a proxy for interspecific competition, we used the same species traits relevant as Gillmann et al. (2024a): feeding type (competition for food), habitat preference (microscale space competition), and stream zonation preference (macroscale space competition). We obtained trait values from the freshwaterecology.info database (Schmidt-Kloiber & Hering, 2015), which provides trait affinities on a 10-point scale. Competition was defined as the proportion of trait overlap between species, indicating similar habitat and food preferences. Using the Gower Similarity from the “gawdis” package in R (v.0.1.5, de Bello et al., 2021), we calculated the trait similarities for each taxa pair per site and year. A lower Gower distance indicates higher similarity and thus higher likelihood of competition. We then summarized these values to the mean Gower similarity per site and year, which we refer as ‘competition’ in the following.

### 2.2 Abiotic data

#### 2.2.1 Catchment land use

For each sampling site, we quantified the catchment land use following the methodology used by Schürings et al. (2024 b, c). Upstream catchments for Germany-wide monitoring sites were delineated using ESRI ArcView 3.3 with a 10 m resolution Digital Elevation Model (DEM) and visually verified. Catchments for the Boye dataset (Gillmann et al., 2024a) were delineated using SWAT+ (Bieger et al., 2017), based on a 10 m DEM of the EGLV (https://www.eglv.de/en/). Agricultural land use data for 2017, derived from random forest classification of Sentinel-2, Landsat 8, and Sentinel-1 data by Blickensdörfer et al. (2022), were used to quantify the percentage cover of cropland within a catchment using ESRI ArcGIS Pro 2.9.0 and Python 3.7. Urban areas and forests for 2016 were included using data from Griffiths et al. (2019).

The temporal mismatch between crop maps (2016-2017) and biological data (2004–2022) was averaged out by the sheer size dataset (1568 sites). Additionally, while crop rotations vary annually, cropland percentages remain stable on larger scales and Schürings et al. (2024b) found only minor differences when relating land use from different years with biota. Extreme weather responses, such as those to 2018’s drought period (Blickensdörfer et al., 2022), were considered by using the ‘year of biological data’ as a random factor in our models.

#### 2.2.2 Physio-chemical parameters

Alongside the macroinvertebrate sampling, at all sites also physio-chemical parameters were measured between monthly to yearly intervals following a methodology of Schneider et al. (2003). The physio-chemical parameters measured included conductivity, nitrite, nitrate, ammonium, phosphate, oxygen, temperature and pH. We linked the physio-chemical sample timely closest but prior to the biological sample with the macroinvertebrate information. Since the Germany-wide data originated from different federal states, the data were on forehand harmonized and standardized to the same unit. Also, not for all physio-chemical parameters (conductivity, nitrate, nitrogen, ammonium, phosphate, oxygen, temperature and pH) measurements were available at each site (e.g. caused by measurement errors). To address these gaps, missing values were imputed based on the existing parameters using an iterative imputation algorithm based on random forest (missForest) with ntree=100, a method which has been shown to perform well for data gaps extending up to 50% of the values (Stekhoven and Bühlmann, 2012; Tang and Ishwaran, 2017).

#### 2.3 Statistical analysis

First, the overall stress from various abiotic stressors was quantified using principal component analysis (PCA) with all measured physio-chemical parameters, using the R package vegan (Oksanen et al., 2022). The value of the first PCA axis was saved and used in subsequent analyses as an overall measure of abiotic stress. Next, we calculated the changes (delta) in ecological status, abiotic stress, and competition between consecutive time points. For example, if a site was sampled in 2004, 2007, and 2010, we computed two delta values: one for 2007 (the difference to 2004) and one for 2010 (the difference to 2007). These changes were referred to as change in ecological status, abiotic stress and competition. We then split the dataset in positive and negative changes of ecological status (delta ecological status) to create both a recovery and degradation sub-dataset, alongside the full dataset. Note, that a particular site can be assigned to both recovery and degradation, when both increasing and decreasing delta values were calculated.

To investigate the relation between changes in abiotic stress and changes in ecological status, we fitted a generalized linear mixed model (GLMM) with zero inflated gamma distribution and log link, using the “glmmTMB” package (v 1.1.8, Brooks et al., 2017) and calculated a pseudo-R^2^ (from here on referred to as R^2^). We used the change in ecological status as response and the change in abiotic stress (PC axis 1) as predictor, while river type (Pottgiesser and Sommerhäuser, 2008) as well as date of biological sampling were used as random factors. Residuals suggested no temporal autocorrelation and no spatial autocorrelation was assumed, given that all sites have distinct sub-catchments and previous studies using the same sampling sites did not find strong residual autocorrelation (e.g. Schürings et al., 2024b). To better account higher density in the center and the large heterogeneity, we also set up a weighted LMM, by increasing the weight of the outliers by squaring the predicted residuals as a function of the response. To compare this relation between recovery and degradation conditions (research question 1), we also fitted the same GLMMs using the recovery and degradation sub-datasets (i.e. those time series that improved and declined in ecological status). To generate the needed ranges of response variables between 0 and 1 for the different models, we transformed the entire dataset using the formula y = (y+1)/2 and multiplied the values of the degradation sub-dataset (ranging between -1 and 0) by -1. For the comparison of the regression coefficients between recovery and degradation datasets, we inversed the regression coefficient of the degradation sub-dataset.

To investigate the relation between changes in competition and in ecological status for recovery and degradation conditions (research question 2), we fitted a similar generalized linear mixed model (GLMM) with zero inflated gamma distribution and both a weighted LMM for the full model and GLMM for the full model. Additionally, we set up the same GLMMs for both the recovery and degradation sub-dataset. In all models, we used the change in ecological status as response, the change in competition as predictor and river type as well as date of biological sampling as random factors. We applied the same transformation for the degradation sub-dataset and compared the regression coefficients between recovery and degradation after inversing the regression coefficient of the degradation sub-dataset.

To investigate the mediating influence of other variables on the recovery and degradation potential of rivers (research question 3), we set up additional GLMs (with zero inflated gamma distribution) for the full dataset and the recovery and degradation sub-datasets, using the change in ecological status as response and the different variables catchment land use, river systems, river type, date of biological sampling, and category (whether the river is assigned as ‘natural’ or ‘heavily modified’) as predictor. We additionally applied spearmen rank correlation between the change in ecological status and the percentage cover of different land use categories.

## 3. Results

The directionality of most prevailing abiotic stressors i.e., nutrients and conductivity were positioned along PC axis 1 (Figure 1). The change in temperature was opposing change in oxygen content, and both were strongly related. to PC axis 2. The PC axis 1 therefore mainly represented the intensity of nutrients and conductivity and related negatively to the ecological status.

**Figure 1:**
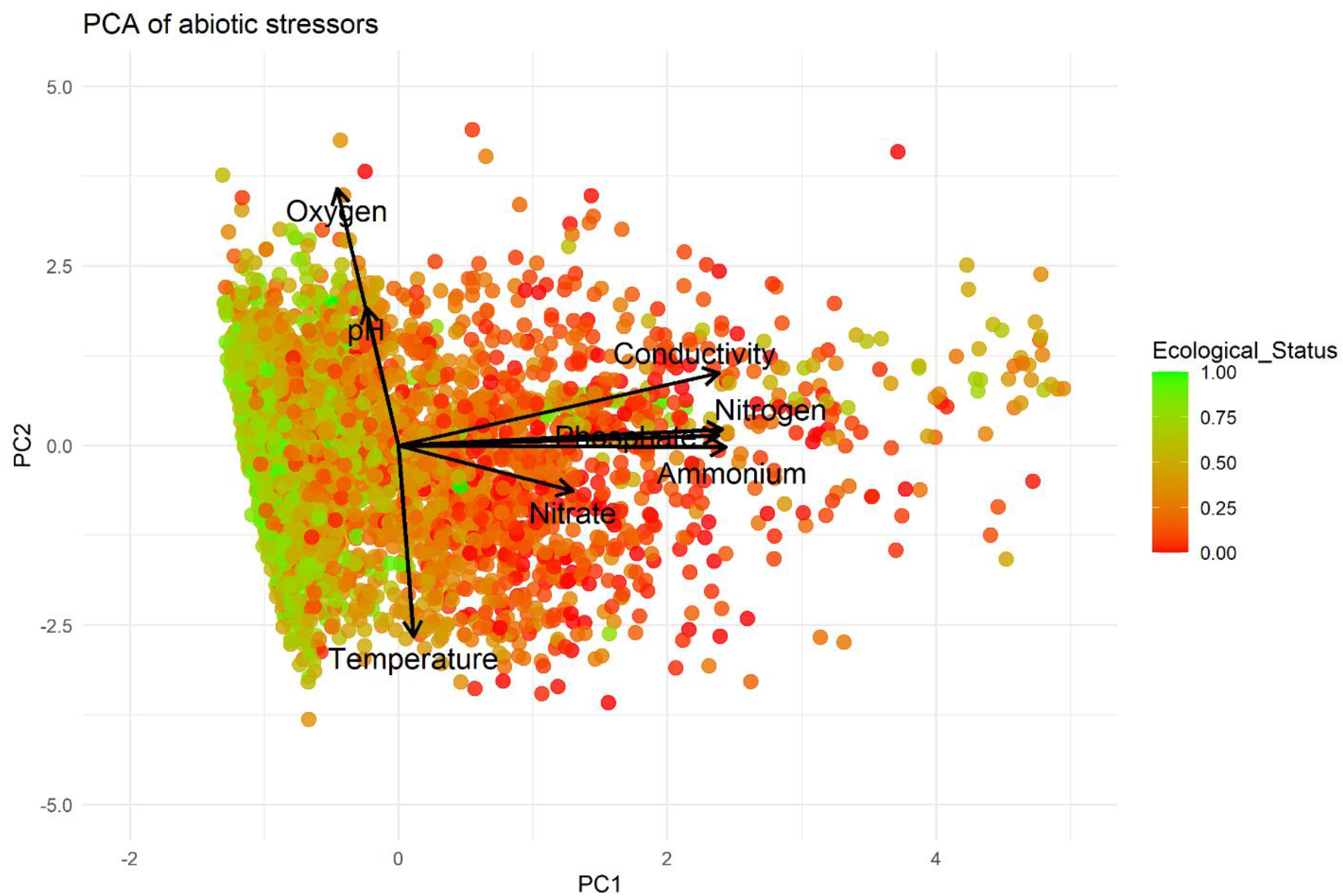
Principal Component Analysis (PCA) of the different abiotic parameters Oxygen, pH, Temperature, Conductivity, Phosphate, Nitrogen, Ammonium and Nitrate. The individual sites are colored based on the ecological status measure.

### 3.1 The relationship between change in abiotic stress and ecological status is similar in recovery and degradation conditions

A negative relation between the change in ecological status and the change in abiotic stress was observed with R^2^ = 0.14 for the unweighted LMM and 0.39 for the weighted LMM (Figure 2a). There was, however, no major difference in the strength of the relation between the changes in abiotic stress and ecological status when differentiating between recovery and degradation conditions (Figure 3a). If assessed individually, the regression coefficient was -0.1 for both conditions.

**Figure 2:**
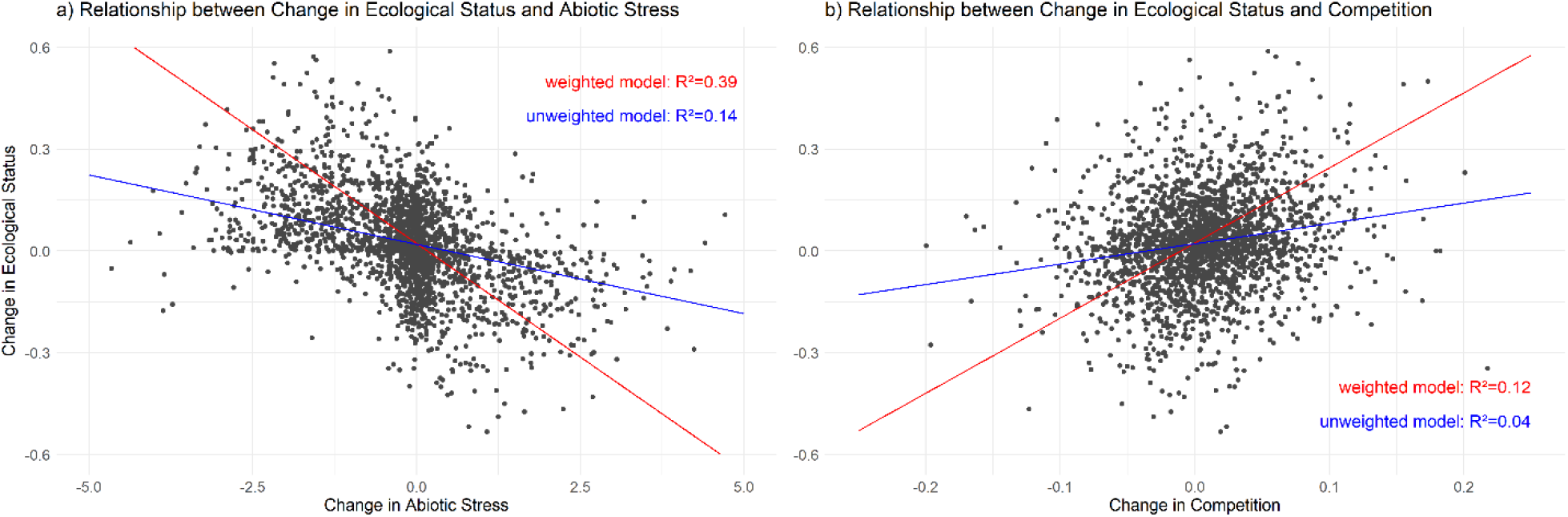
LMMs (blue line) and weighted LMMs (red line) with the change in Ecological status and change in Abiotic Stress (left) and the change in Ecological status and change in Competition (right). Each model has 1 predictor and date of biological sampling as well as river types as random factors.

**Figure 3:**
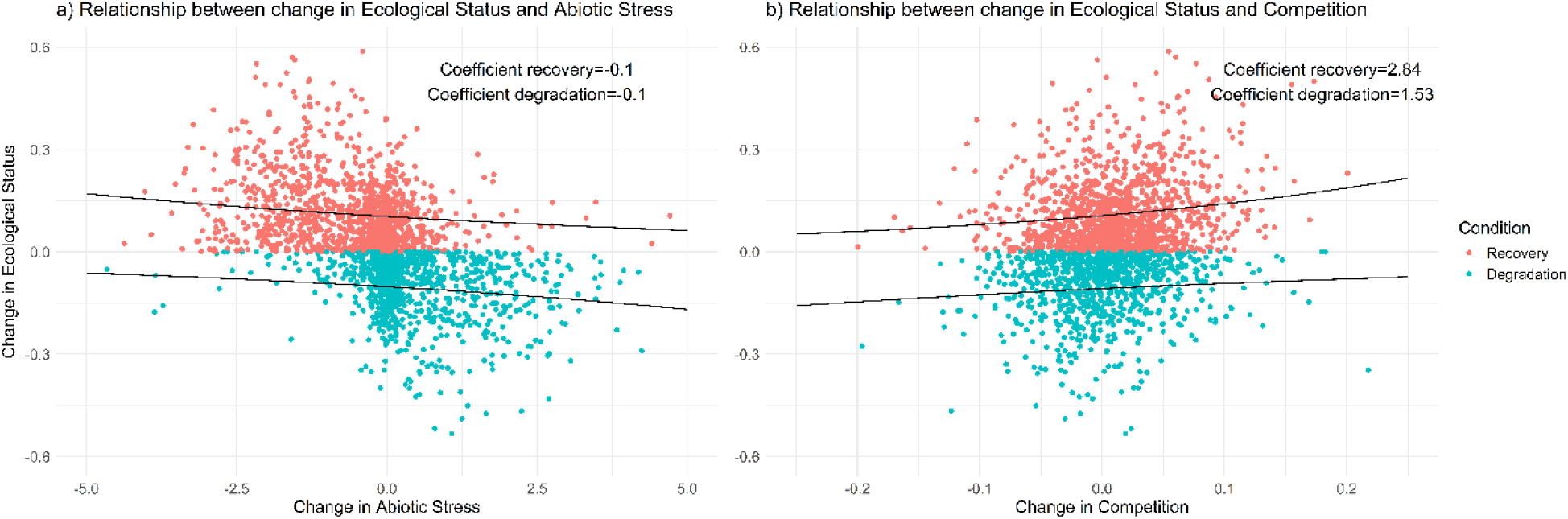
GLMMs with zero inflated Gamma distribution and log link for two sub-datasets of recovery (red) and degradation conditions (blue) with the change in Ecological status as response and in a) change in Abiotic Stress and predictor and b) change in Competition as predictor. River type and date of biological sample are random factors.

### 3.2 The relationship between the change in competition and ecological status is stronger under recovery conditions compared to degradation conditions

In the complete dataset, the correlation between changes in ecological status and competition is slightly positive, with an R^2^ value of 0.04 for the unweighted LMM and 0.12 for the weighted LMM (Figure 2b). When separating recovery and degradation sites and using GLMMs (Figure 3b), a positive relationship is observed for both. Notably, the regression coefficient is nearly twice as high in recovery sites (2.84) compared to degradation sites (1.53).

### 3.3 Land use and river types mediate recovery and degradation potential

The results of the GLMs (Table 1) showed no influence of the additional variables when assessing the full model (no significant variables and R^2^ < 0.02). While for river category (HMWB, NWB) and date, there was also no relation when separating recovery and degradation sites, river systems might have a minor influence on degradation sites. River types and land use both showed small significant influences when separating between recovery and degradation sites (R^2^ up to 0.05).

**Table 1:** Model outputs from GLM with zero inflated gamma distribution. The change in ecological status was used as response and each of the different variables as predictor. Shown are the R^2^ and the significance level p. When p< 0.05 the values are presented in bold.

|  | River systems | River status | Date | River types | Land use |
| --- | --- | --- | --- | --- | --- |
| Full dataset | $R^2 = 0.01$<br>$p > 0.05$ | $R^2 < 0.01$<br>$p > 0.05$ | $R^2 < 0.01$<br>$p > 0.05$ | $R^2 = 0.01$<br>$p > 0.05$ | $R^2 < 0.01$<br>$p > 0.05$ |
| Recovery sub-dataset | $R^2 < 0.01$<br>$p > 0.05$ | $R^2 < 0.01$<br>$p > 0.05$ | $R^2 < 0.01$<br>$p > 0.05$ | **$R^2 = 0.05$**<br>**$p < 0.001$** | **$R^2 = 0.01$**<br>**$p = 0.04$** |
| Degradation sub-dataset | **$R^2 = 0.01$**<br>**$p = 0.008$** | $R^2 < 0.01$<br>$p > 0.05$ | $R^2 < 0.01$<br>$p > 0.05$ | **$R^2 = 0.05$**<br>**$p < 0.001$** | **$R^2 = 0.02$**<br>**$p = 0.01$** |

The influences for river systems and river types were unclear (no clear trend could be found), wherefore, the influence of river systems was not further investigated, and the river types were only accounted for in form of random factors in the GLMMs for part 3.1 and 3.2. When relating the percentage of the different catchment land use categories - forest, urban areas and cropland - with the change in ecological status in recovery and degradation sites using spearman rank correlation, a small difference was observed (Table 2). While the change in ecological status related negatively to forest cover but positively to cropland and urban catchment land use, when using the recovery sub-dataset, the relation was contrary in the degradation sub-dataset. Here, forest land use related positively, while cropland and urban areas related negatively.

**Table 2:** Spearman rank correlation between the change in ecological status and the percentage catchment cover of forest, cropland and urban areas, when using the full dataset, and the recovery as well as degradation sub-datasets.

|  | Forest | Cropland | Urban |
| --- | --- | --- | --- |
| Full dataset | $\rho = -0.03$ | $\rho = 0.02$ | $\rho = 0.03$ |
| Recovery sub-dataset | $\rho = -0.05$ | $\rho = 0.07$ | $\rho = 0.05$ |
| Degradation sub-dataset | $\rho = 0.09$ | $\rho = -0.07$ | $\rho = -0.11$ |

## 4. Discussion

This study is the first large-scale study investigating the Asymmetric Response Concept (ARC) of Vos et al. (2023), particularly the two parts of the hypothesis of species tolerances and interspecific competitions are analyzed. Change in abiotic stress appears to be a major cause for both degradation and restoration, while competition is much less important but appears to be more relevant in restoration scenarios compared to degradation.

### 4.1 Abiotic stress and the Ecological status

As expected and observed by a multitude of other studies (e.g Johnson et al., 2013; Wagenhoff et al., 2012), we found a clear relationship between the change in ecological status and the change in abiotic stress, mainly nutrient concentrations and conductivity. Following the Asymmetric Response Concept (Vos et al., 2023), we expected a stronger relationship between the change in abiotic stress and ecological status (as a proxy for overall stress tolerances) in phases of degradation as compared to recovery conditions, the strength of relation appeared similar (both regression coefficients of -0.1). However, given that the relationship for the split datasets was relatively small this should be taken with care.

### 4.2 Competition

The relationship between changes in ecological status and changes in trait similarity was found to be slightly positive overall. This suggests that if the ecological status improves, there tends to be a higher overlap in trait affinities among species, indicating increased interspecific competition. Notably, this positive relationship between changes in ecological status and competition was stronger during recovery conditions compared to degradation conditions.

Although the relationship between changes in abiotic stress and ecological status showed similar strength in both recovery and degradation conditions, the more pronounced positive relation between improved ecological status and increased competition during recovery might still contribute to the complexity of the recovery process, suggesting that in recovery conditions species compete more intensively for the same resources. Thus, the stronger positive relationship between ecological status improvement and competition during recovery conditions could be a relevant factor in influencing the dynamics of recovery, even when reductions in abiotic stress are achieved, which is in line with the ARC (Vos et al., 2023).

### 4.3 Other influencing variables

When assessing the entire dataset, the other potentially influencing variables (river systems, river status, date, river types and land use) showed no relation to the change in ecological status. However, when distinguishing between recovery and degradation sites, small trends could be observed. While for river types, the direction was unclear, it was used as a random factor in the analysis (as has been done in previous studies to account for variables not originating from the ones considered (e.g. Schürings et al., 2024b).

Even though the correlations were small (Spearman ρ between -0.05 and 0.11), we observed a trend: degradation (a decrease in ecological status) is more apparent in less intensely used catchments, such as those covered by forests, while recovery is more likely in intensely used catchments, such as cropland or urban areas. This underlines the relationship of biota to land use intensity (e.g., Chen et al., 2023; Schürings et al., 2024a). In intensively used catchments, the ecological status is already low, making further deterioration less likely than recovery. Conversely, in less intensely used catchments, the higher initial ecological status makes them more susceptible to degradation.

### 4.4 Methodological Reflections

The relationships found in this study are small in magnitude and should be considered associations rather than direct causal effects. Recovery and degradation sites were categorized based on changes in ecological status between two time points, meaning recovery sites do not necessarily represent recently restored reaches. This complicates to unravel whether recovery is slowed down by interspecific competition, making the observed relationships more tentative. The datasets used have some limitations. Abiotic parameters were measured as spot measurements, which may not capture the full environmental variability. Additionally, several potentially relevant stressors, such as hydromorphological degradation (Lorenz et al., 2009) and micropollutants (Markert et al., 2024) were not considered. Using trait overlap as a proxy for competition is a broad measure and may not fully capture the complexity of competitive interactions among species. These considerations suggest the need for cautious interpretation of the results and indicate that future research should aim to address these methodological aspects to better understand the variables influencing ecological recovery and degradation.

### 4.5 Study implications

This study provides support for the hypotheses of the ARC (Vos et al., 2023), emphasizing the importance of context in ecological recovery and degradation processes. These findings underscore that recovery and degradation are context specific and likely mediated by species tolerances and interspecific competition.

Even when using crude proxies, this study supports the conjecture that changes in water quality are a primary driver of ecological status. This suggests that improvements in water quality can lead to significant ecological benefits. However, more localized stressors and future studies would benefit from including stressors such as pesticides and emergent pollutants (Lencioni et al., 2020; Link et al., 2022; Schäfer, 2019) and changes in hydromorphological conditions (Lorenz et al., 2009; Verdonschot et al., 2016), which also play crucial roles in shaping ecological communities. Overall, these insights contribute valuable knowledge to the field of freshwater ecology, emphasizing the need to consider both broad-scale water quality improvements and local-scale stressors in restoration and conservation efforts.

## Acknowledgements

This paper results from the Collaborative Research Centre 1439 RESIST (Multilevel Response to Stressor Increase and Decrease in Stream Ecosystems; www.sfb-resist.de) funded by the Deutsche Forschungsgemeinschaft (DFG, German Research Foundation; CRC 1439/1, project number: 426547801).

